# At-home saliva sampling in healthy adults using CandyCollect, a lollipop-inspired device

**DOI:** 10.1101/2023.01.14.524039

**Authors:** Wan-chen Tu, Anika M. McManamen, Xiaojing Su, Ingrid Jeacopello, Meg G. Takezawa, Damielle L. Hieber, Grant W. Hassan, Ulri N. Lee, Eden V. Anana, Mason P. Locknane, Molly W. Stephenson, Victoria A. M. Shinkawa, Ellen R. Wald, Gregory P. DeMuri, Karen Adams, Erwin Berthier, Sanitta Thongpang, Ashleigh B. Theberge

**Author notes:** **Corresponding Author Ashleigh B. Theberge** − Department of Chemistry, University of Washington, Seattle, Washington 98195, United States; Department of Urology, School of Medicine, University of Washington, Seattle, Washington 98195, United States;, **Sanitta Thongpang** − Department of Chemistry, University of Washington, Seattle, Washington 98195, United States; Department of Biomedical Engineering, Faculty of Engineering, Mahidol University, Nakorn Pathom 73170, Thailand. These authors contributed equally to this work (Wan-chen Tu, Anika M. McManamen).

## Abstract

Respiratory infections are common in children, and there is a need for user-friendly collection methods. Here, we performed the first human subjects study using the CandyCollect device, a lollipop inspired saliva collection device.^1^ We showed the CandyCollect device can be used to collect salivary bacteria from healthy adults using *Streptococcus mutans* and *Staphylococcus aureus* as proof-of-concept commensal bacteria. We enrolled healthy adults in a nationwide (USA) remote study in which participants were sent study packages containing CandyCollect devices and traditional commercially available oral swabs and spit tubes. Participants sampled themselves at home, completed usability and user preference surveys, and mailed the samples back to our laboratory for analysis by qPCR. Our results showed that for participants in which a given bacterium (*S. mutans* or *S. aureus*) was detected in one or both of the commercially available methods (oral swab and/or spit tubes), CandyCollect devices had a 100% concordance with the positive result (n=14 participants). Furthermore, the CandyCollect device was ranked the highest preference sampling method among the three sampling methods by 26 participants surveyed (combining survey results across two enrollment groups). We also showed that the CandyCollect device has a shelf life of up to 1 year at room temperature, a storage period that is convenient for clinics or patients to keep the CandyCollect device and use it any time. Taken together, we have demonstrated that the CandyCollect is a user-friendly saliva collection tool that has the potential to be incorporated into diagnostic assays in clinic visits and telemedicine.

**For Table of Contents Only:** 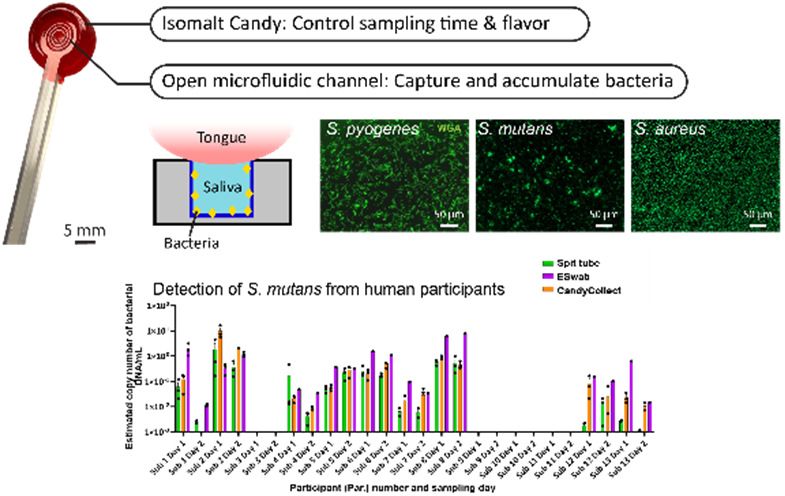

## INTRODUCTION

Infectious respiratory pathogens are a major health challenge worldwide. Children, in particular, are frequently affected by respiratory diseases.^2^ The Covid-19 pandemic has illustrated the importance of global pandemic preparedness, and in particular the need to develop more comfortable and user-friendly sampling methods to test for pathogens. We developed a lollipop-inspired saliva collection device called CandyCollect to enable user-friendly sampling in both children and adults (Figure 1). Here, we conducted a human subjects study as a proof-of-concept to demonstrate functionality of the CandyCollect device for capturing bacteria from healthy adults and evaluate the comfort and user experience in comparison to standard saliva collection methods.

**Figure 1.**
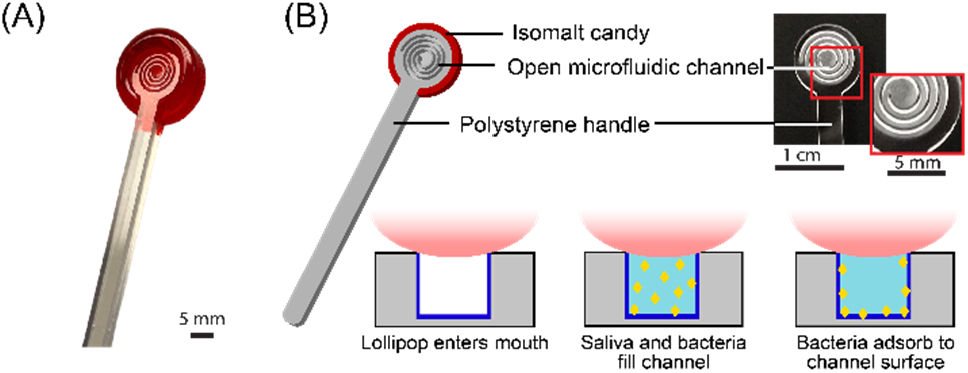
(A) The photo of the CandyCollect device. The CandyCollect device is composed of a polystyrene stick with a microfluidic channel and red isomalt candy. The open-fluidic channel is designed to prevent the tongue from removing the collected bacteria, also accumulating bacteria during the sampling time. The candy flavoring functions as a built-in timer for sampling time (i.e., dissolving time of the candy). (B) This figure is reproduced from Lee et al. ^1^ (Figure 1B) with permission from the Royal Society of Chemistry.

As previously reported, the CandyCollect is a saliva sampling device designed to collect *Streptococcus pyogenes* for the diagnosis of Group A streptococcus (GAS) pharyngitis, commonly referred to as strep throat.^1^ Strep throat is most commonly seen in children.^3^ It is easily treatable with antibiotics when diagnosed, however diagnosis can be thwarted by invasive sampling methods which discourage children (and adults) from successfully completing the sampling process and may result in decreased yields.^4^ The gold standard method for diagnosing strep throat is a pharyngeal swab coupled with bacterial culture,^5,6^ however, qPCR has become a new tool that can be implemented in diagnosis of strep throat, allowing saliva sampling as a means for diagnosis.^7,8^ Current methods for saliva sampling include spit tubes (e.g., SpeciMAX Stabilized Saliva Collection Kit), drooling (e.g., SalivaBio Saliva Collection Aid), swabs (e.g., Eswab™) and cotton rolls (e.g., Salivette^®^, SalivaBio Oral Swab). In recent years, others have also developed lollipop-inspired devices, such as Self-LolliSponge™ (with lemon-aromatized cap), and non-conventional sampling devices using absorbing materials, e.g. V-Chek™ test card and Whistling™ midstream test.^9-13^ The CandyCollect device was designed to facilitate easy, non-invasive, at-home sampling of saliva, particularly for children, which is then shipped to a lab for analysis and diagnosis.^1^ The device sampling mimics the action of eating a lollipop, featuring a polystyrene stick with an open microfluidic channel for bacterial capture and isomalt candy that functions as a timer to ensure sufficient sampling time.

Here we demonstrate the versatility and functionality of the CandyCollect device through experimentation and at-home human research studies using commensal bacteria *Streptococcus mutans* and *Staphylococcus aureus* for proof-of-concept. Commensal bacteria exist in the microbiome of healthy hosts.^14,15^ Using commensal bacteria as our analytes of interest, as opposed to *S. pyogenes*, which we reported previously,^1^ allowed us to enroll healthy participants for our human subjects research. In doing this, our target population was much broader than the limited population of those with strep throat, allowing greater diversity and easier enrollment of participants. This also provided the opportunity to test different bacteria and access the versatility of CandyCollect devices as a broader microbial sampling method. *S. mutans* and *S. aureus* were specifically selected as analytes due to their high prevalence in the healthy adult population.^16-19^ The prevalence of *S. mutans* and *S. aureus* has been reported to vary between 80-87% and 18-39% in healthy adults, respectively.^16-19^ In-lab experiments demonstrated the CandyCollect device can capture these commensal bacteria, and, through elution and qPCR, quantify their concentration. Our human subjects study established two key findings regarding the CandyCollect device: (1) the CandyCollect device was the preferred sampling method among participants compared to conventional sampling methods, and (2) the CandyCollect device captures *S. mutans* and *S. aureus*.

## METHODS

### CandyCollect devices fabrication

CandyCollect device stick fabrication: The CandyCollect devices were milled out of 2 mm and 4 mm polystyrene sheets (Goodfellow, Cat# 235-756-86 and 700-272-86, respectively) using a DATRON computer numerically controlled (CNC) milling machine (Datron) (Figure S1). Devices were then sonicated in isopropanol (IPA) (FisherScientific, A451-4) and 70% v/v ethanol (FisherScientific, Decon™ Labs, 07-678-004).

Plasma treatment of CandyCollect devices: Devices were plasma treated with oxygen using the Zepto LC PC Plasma Treater (Diener Electronic GmbH, Ebhausen, Germany). The protocol for plasma treatment is consistent with our previous publication, but in brief, gas was removed from the chamber down to a pressure of 0.20 mbar, oxygen gas was supplied up to 0.25 mbar for 2 minutes and then a 70 W voltage was applied for 5 minutes.^1^ Following plasma treatment, devices for spike sample experimentation were ready for use.

Preparation of CandyCollect devices for human subjects study: Candy was applied to CandyCollect sticks in a kitchen following the hygiene guidance outlined in the Washington State Cottage Food Operations Law (RCW 69.22.040(2b-f(ii-iv))). Lab members who prepared CandyCollect were trained in food safety, had a Food Worker Card (WA State), and wore gloves and a mask during food preparation. The isomalt candy was prepared as described in our previous paper.^1^ In brief, isomalt was gradually added to water. Food coloring was added with the last portion of isomalt. Once dissolved the isomalt was then heated to either 171 °F (dissolve time <20 min) or 165 °F (dissolve time >20 min). Once target temperature is reached, the pot containing isomalt was quickly placed in room temperature water to initiate cooling. At this time strawberry candy flavoring was quickly added to the mixture, and the isomalt poured onto a marble slab to set. After plasma treatment, CandyCollect polystyrene sticks for the human subjects study were cleaned using hot water and dish soap. Small portions of the isomalt candy were remelted and applied to the CandyCollect sticks using a silicone mold. Once the candy was applied to the sticks, the candy was cooled, the device mass was recorded, and the CandyCollect devices were placed into polypropylene bags and heat sealed. Devices were stored in food preparation containers with a desiccant (Amazon, Cat# B00DYKTS9C) until being sent to participants. CandyCollect devices used in this study had masses ranging from 1.2-1.9 grams on both days (Table S1-2).

### Bacteria culture

Liquid media preparation: The *S. mutans* culture media (trypticase soy yeast extract medium) was prepared based on the method from DSMZ website.^20^ The *S. aureus* culture media (Tryptic Soy Agar/Broth) was prepared based on the protocol from ATCC.^21^ The *S. pyogenes* culture media was prepared following the protocol from Gera & McIver, 2013.^22^ All liquid media were autoclaved at 121 °C for 30 min, cooled to room temperature, and stored at 4 °C.

Agar plate preparation: 7.5g agar (BD Difco™ Dehydrated Culture Media: Potato Dextrose Agar, Fisher Scientific, Cat# DF0013-17-6) was added to the 500 mL of liquid media, then autoclaved at 121 °C for 30 min. 15 mL of liquid media with agar was added to petri dishes, left to cool overnight, and stored at 4°C until needed for.

*S. mutans, S. aureus*, and *S. pyogenes* maintenance in agar plate: *S. mutans* was prepared from *Streptococcus mutans* Clarke (American Type Culture Collection, ATCC^®^, Cat# 25175™). 880 μL of liquid media was added to freeze-dried *S. mutans*, and the bacteria suspension was transferred into a 10 mL tube. Additional liquid media was added for a total volume of 8.8 mL. *S. aureus* was prepared from *Staphylococcus aureus* subsp. aureus Rosenbach (American Type Culture Collection, ATCC^®^, Cat# 25923™). Freeze-dried *S. aureus* was rehydrated in 910 μL of liquid media. *S. pyogenes* was prepared from *Streptococcus pyogenes* Rosenbach (American Type Culture Collection, ATCC^®^, Cat# 700294™). Freeze-dried *S. pyogenes* was rehydrated with 1 mL liquid media, and then transferred to another conical tube containing 4.4 mL of liquid media. To maintain the bacteria, *S. mutans, S. aureus*, and *S. pyogenes* were streaked on their own agar plates by sterile disposable inoculating loops (Globe Scientific, Fisher Scientific, Cat# 22-170-201). The agar plates were incubated at 37 °C with 5% carbon dioxide overnight, then stored at 4°C until needed for experimentation.

### In-lab capture of bacteria

Incubation of *S. mutans, S. aureus*, and *S. pyogenes* in liquid media: To ensure a pure culture, fresh *S. mutans, S. aureus*, and *S. pyogenes* from agar plates were inoculated in liquid media and cultured at 37 °C with 5% carbon dioxide in the incubator one day prior to an experiment.

Capturing, fixing, and staining bacteria: The procedures for capturing, fixing, and staining bacteria in liquid media are detailed in our previous paper.^1^ In brief, after culturing overnight, the bacteria suspensions were homogenized with vortexing and added to each CandyCollect device at a volume of 50 μL (devices negative controls were loaded with 50 μL of PBS). Bacteria were incubated in the device for 10 min. For devices that were imaged, bacteria were fixed with 4% paraformaldehyde (PFA) for 15 min, and 50 μL of Alexa Fluor™ 488 Wheat Germ Agglutinin (WGA, Invitrogen™, Fisher Scientific, Cat# W11261, 1 mg/mL) at 1:500 dilution (v/v) was added to the channel for staining *S. aureus* and *S. pyogenes*; 50 μL of 1:200 (v/v) WGA was added for staining *S. mutans*. An additional three devices were evaluated for a mixture of *S. mutans, S. aureus* and *S. pyogenes*, each at a concentration of 10^4^ CFU/mL to match physiological bacterial concentration for detection of bacteria in a mixture.

### Fluorescence imaging and quantification

Fluorescent images of *S. mutans, S. aureus*, and *S. pyogenes* were obtained on a Zeiss Axiovert 200 with a 10× (0.30 NA) objective coupled with Axiocam 503 mono camera (Carl Zeiss AG, Oberkochen, Germany). Four regions of interest were randomly chosen from each device to avoid bias from any regions. The contrast was adjusted uniformly and integrated densities of three regions of interest from each image were quantified using Fiji (ImageJ) software. The details about imaging and quantification for both bacteria followed the protocol from the previous paper.^1^

### Elution of *S. mutans, S. aureus*, and *S. pyogenes* from CandyCollect devices

The buffer used to elute bacteria captured on CandyCollect devices was phosphate buffered saline (PBS) (Gibco™, Cat# 10010023) with 5% Proteinase K (Thermo Scientific™, Cat# EO0491). 300 μL elution buffer and 100 μL of 0.1 mm Zirconia/Silica beads (BioSpec Products, Cat# 11079101Z) were added in 14 mL round bottom tubes (Corning, Falcon^®^, 352001) containing CandyCollect devices. After incubating the tubes at 37 °C for 10 min and vortexing for 50 s, CandyCollect devices were left in the elution buffer at 4 °C for 90 min. The bacteria suspension and beads were then transferred from the 14 mL round bottom tubes to 2 mL screw cap microtubes (ThermoFisher, Cat# 3490). The samples were beat-beaten in a MiniBeadBeater (BioSpec Products, Bartlesville, OK USA), and stored at -20 °C before analysis.

Additional elution buffers evaluated included (1) ESwab™ buffer (Becton, Dickinson and Company, Cat# R723482) with 5% Proteinase K; (2) ESwab™ buffer with 5% ethanol; (3) phosphate buffered saline (PBS) with 2% SDS (sodium dodecyl sulfate); and (4) ESwab™ buffer with 2% SDS. The elution procedures were the same as mentioned above.

### Isolation, purification, and enrichment of genomic DNA from *S. mutans, S. aureus*, and *S. pyogenes*

DNA was isolated from bacterial lysates using the Mag-MAX™ Total Nucleic Acid Isolation Kit (ThermoFisher Scientific, Cat# AM1840) according to the “Purify the nucleic acid” protocol supplied by the manufacturer. In brief, 115 μL of sample was added to the provided processing plate. 60 μL of 100% IPA was added to each well containing a sample and the plate was shaken for 1 min. 20 μL of bead mix was then added to each well, and the plate was shaken for 5 min to allow DNA to bind to the beads. Beads were captured using a magnetic 96-well separator (Thermofisher, Cat# A14179) and supernatant was discarded. Four washes (two using Wash Solution 1 and additional two using Wash Solution 2 provided by the kit) were performed with shaking for 1 min each and supernatant was discarded between each wash. After final wash, beads were dried and then 23 μL of 65°C elution buffer was added to each sample to elute DNA from the beads. By using these methods, DNA was five-fold concentrated compared to the unprocessed bacterial lysates. The purified bacterial genomic DNA was used as a template in the qPCR assay.

### Quantitative PCR assay for detection of *S. mutans, S. aureus*, and *S. pyogenes*

The species-specific genes, *S. mutans gtfB* (accession number M17361), encoding glucosyltransferases, and *S. aureus nuc* (accession number CP000046), encoding a thermonuclease, were used for qPCR detection of *S. mutans* and *S. aureus*, respectively. The primers/probe sequences for *gtfB* were adopted from Lochman et al., 2020,^23^ the forward primer: 5’-CCT ACA GCT CAG AGA TGC TAT-3’; the reverse primer: 5’-GCC ATA CAC CAC TCA TGA ATT-3’; the probe: 5’-/56-FAM/TGG AAA TGA/ ZEN/CGG TCG CCG TTA T/3IABkFQ/ -3’. Primers/probe sequences for *nuc* were adopted from Wood et al., 2021 and Galia et al., 2019,^24, 25^ with minor modifications to both forward and reverse primers, the forward primer (F1): 5’-GGC ATA TGT ATG GCA ATC GTT TC-3’; the reverse primer (R1): 5’-CGT ATT GTT CTT TCG AAA CAT T-3’; the probe sequence: 5’-/56-FAM/ATT ACT TAT AGG GAT GGC TAT C/3MGB-NFQ/ -3’. The modified primers were accessed for their specificity using NCBI Blast tool and verified by qPCR assay with purified DNA from *S. aureus*. Details can be found in Table S3 and Figure S2-3 and the discussion in the SI. All primers and probes for *S. mutans* and *S. aureus* were ordered from IDT (Integrated DNA Technologies, Inc., Coralville, IA, USA). qPCR was performed using PerfeCTa^®^ qPCR ToughMix (VWR, Cat# 97065-954). The 25 μL reaction volume included 5 μL of DNA template and 20 μL PerfeCTa^®^ qPCR ToughMix with primers/probe in the qPCR assay. For *S. mutans* and *S. aureus* analysis, the final concentrations of both forward and reverse primers were 300 nM and 500 nM, respectively; the probe concentration for both bacteria was 250 nM. The details for the qPCR assay for *S. pyogenes* followed the protocol from our previous paper.^1^ Briefly, the primers/probe sequences for *spy1258* qPCR detection of *S. pyogenes* in our assay were: the forward primer: 5’-GCA CTC GCT ACT ATT TCT TAC CTC AA-3’; the reverse primer: 5’-GTC ACA ATG TCT TGG AAA CCA GTA AT-3’; the probe sequence: 5’-FAM-CCG CAA C”T”C ATC AAG GAT TTC TGT TAC CA-3’-SpC6, “T” = BHQ1.1 For *S. pyogenes*, the primers were ordered from IDT, the probe was ordered from MilliporeSigma.^1^ The 25 μL reaction volume included 10 μL of DNA template and 15 μL PerfeCTa^®^ qPCR ToughMix with primers/probe in the qPCR assay. The final concentrations of both forward and reverse primers were 300 nM; the probe concentration was 100 nM. To quantify the DNA concentrations of samples, 1:10 serial dilution of purified genomic DNA ranging from 25 ng to 25 fg were used as standards for each plate. Each concentration of the standards was allotted into multiple 20 μL aliquots and stored at -80 °C (Figure S4), to ensure the same standards were used for all human subjects samples. No-template controls (NTC) for qPCR and device negative controls (see “in-lab capturing of bacteria”) were also added to the plates. Amplification and detection were performed in 96-well PCR plates using CFX connect Real-Time PCR Detection System (Bio-Rad Laboratories, Hercules, CA, USA) in technical duplicate using the following protocol: 95 °C for 5 min followed by 40 cycles of 15 s at 95 °C and 30 s at 60 °C. The samples were considered positive when the Cq value is within the Cq of the standard curve.

### Human subjects study

Participant characteristics: This study was approved by the University of Washington Institutional Review Board (IRB) under IRB-approved protocol STUDY00013842. Written informed consent was obtained prior to study procedures. A total of 28 healthy volunteers over the age of 18 were recruited using the University of Washington Institute of Translational Health Sciences (ITHS) “participate in research” website along with the study’s website. Inclusion criteria: healthy adults over the age of 18. Exclusion criteria: individuals who are allergic to sugar alcohols or individuals who currently reside in a correctional facility.

### Human subjects’ sample and feedback collection

The entire study was performed remotely using Research Electronic Data Capture (REDCap) for collection of participant information and survey responses (Table S4), and kits were mailed to study participants and returned to the study team by mail (Figure S5). Each kit contained six CandyCollects, six ESwab™ (Becton, Dickinson and Company, Cat# R723482), two SpeciMAX Stabilized Saliva Collection Kit (Spit tube) (Thermo Scientific™, Cat# A50697), and an instruction card. Based on the instruction card, participants collected samples for two days, and followed the same order of collection on both days: first, one SpeciMAX Stabilized Saliva Collection Kit, second, three ESwab™, and third, three CandyCollects. Collection for each method was instructed as follows: for the spit tube, participants spit approximately 1 mL of saliva into tubes provided by SpeciMAX Stabilized Saliva Collection kit; for ESwab™, participants sucked on the swab for 30 s and then kept the swab in the buffer provided by ESwab™; for the CandyCollect devices, participants were asked to suck on the lollipop until the candy was fully dissolved then left each CandyCollect device in an individual empty polypropylene 12 mL round bottom tube (Greiner Bioone, Cat#163261) and record the time required for the candy to dissolve. The samples then were mailed back to our lab for analysis at the end of each day. An electronic survey was sent to each participant on the same day that they collected samples; the survey included CandyCollect dissolving times and any comments they wanted to leave about each of the sampling devices. After the Day 2 survey was completed, a user feedback survey was automatically sent to participants via REDCap. Participants were asked to rank the different sampling methods as well as answer other specific questions related to the CandyCollect device.

We had two rounds of enrollment, targeting 15 participants per group (Figure 2). In group 1, 15 participants were recruited, but one of the participants did not return the kit and was lost to follow-up, so a total of 14 samples were returned and analyzed. Biological data from this group is presented in Figure 4. In group 2, 15 participants were recruited, however one participant was lost to follow-up and did not return their kit. A total of 14 kits were returned. Biological data from these samples is not presented in this paper as the samples are being used in an additional biological investigation.

**Figure 2.**
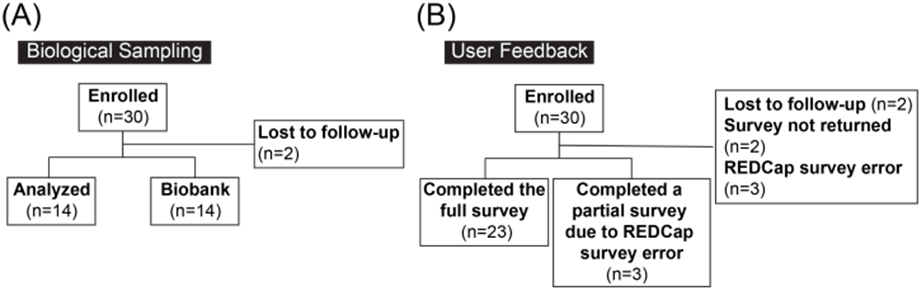
Participant flow diagram. (A) Human subjects samples for biological analysis. (B) User feedback surveys.

**Figure 3.**
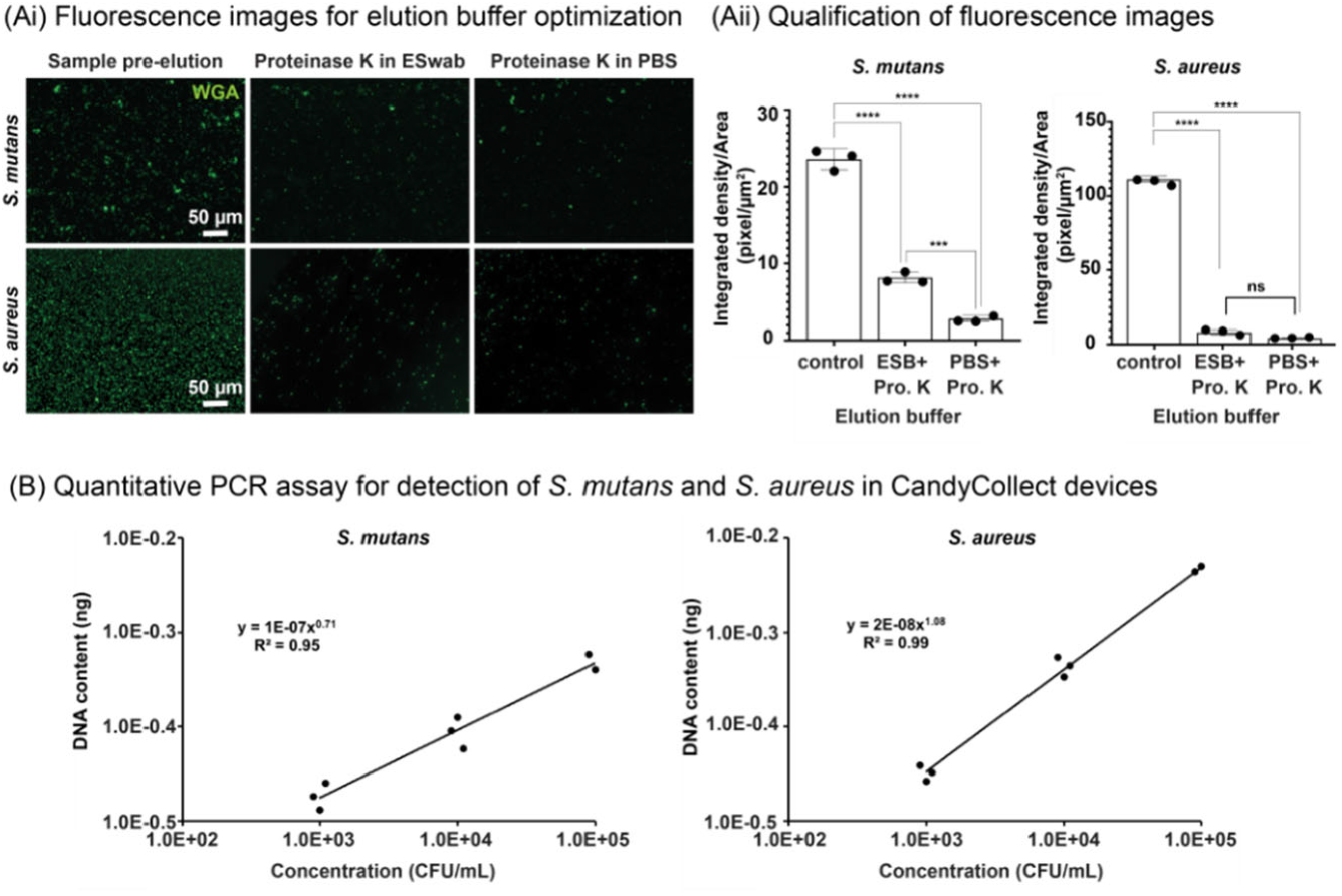
CandyCollect efficiently captures *S. mutans* and *S. aureus* and facilitates quantitative bacterial detection by qPCR. (Ai) Fluorescence microscopy images of *S. mutans* and *S. aureus* before elution (left), and after elution with Proteinase K in ESwab™ buffer (ESB) (middle) and with Proteinase K (Pro. K) in PBS (right). (Aii) Quantification of fluorescence microscopy images. Each data point represents data from one CandyCollect device and is the average of the integrated density per area of 12 (10000 μm^2^) regions of interest (ROI) from 4 images of the CandyCollect device (3 ROIs per image). The bar graph represents the mean ± SEM of n = 3 CandyCollect devices. Data sets were analyzed using one-way ANOVA with Tukey’s multiple comparison test; p-values are indicated for pairwise comparisons: ***p=0.0009, ****p<0.0001. (B) Reported concentration of *S. mutans* (left) and *S. aureus* (right) from qPCR analysis of bacteria eluted from CandyCollect devices using proteinase K in PBS as the CandyCollect elution buffer. Bacteria were incubated in-lab at concentrations of 10^3^, 10^4^ and 10^5^ CFU/mL. *S. mutans* and *S. aureus* were fluorescently labeled with WGA.

**Figure 4.**
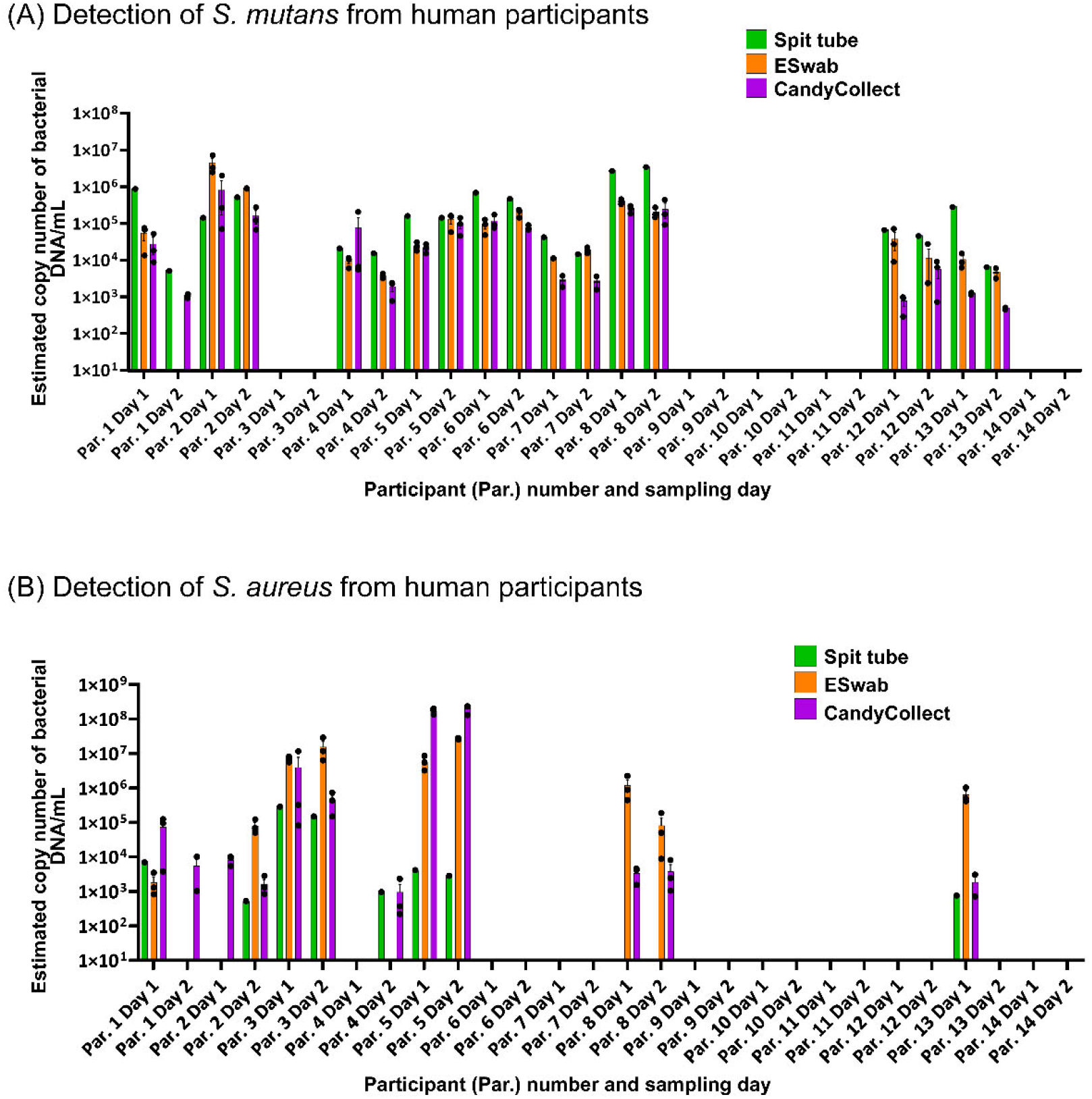
*S. mutans* and *S. aureus* can be captured on CandyCollect devices from all the participants who had positive results from spit tube and/or ESwab™ samples. CandyCollect, ESwab™, and SpeciMAX Stabilized Saliva Collection Kits (Spit tube) were sent to 14 research participants for a proof-of-concept test. The concentrations of (A) *S. mutans* and (B) *S. aureus* from participants’ saliva, collected over two days by three different methods, were analyzed via qPCR and converted to estimated copy number of bacterial DNA/mL in the original saliva sample (see Methods section “Human subject data analysis” for detailed description of calculations). Each dot represents the average of two qPCR technical duplicates from one sample (three samples were collected for the CandyCollect device and ESwab™, and one for the Spit tube each day). Bar graphs represent the mean ± SEM of n = 3 CandyCollect devices or ESwab™. Participants only completed one spit tube per day.

User feedback from groups 1 and 2 is presented in Figure 5. In group 1, all 14 participants answered the survey question corresponding to Figure 5A, however due to an electronic survey error, 3 of the 14 participants were unable to complete the survey questions in Figure 5B. In group 2, 12 of the 14 participants completed the survey questions in Figure 5. This resulted in a total of 26 participants responding to the question in Figure 5A and 23 participants responding to the question in Figure 5B. Protocol for sampling and surveys were identical for both groups.

**Figure 5.**
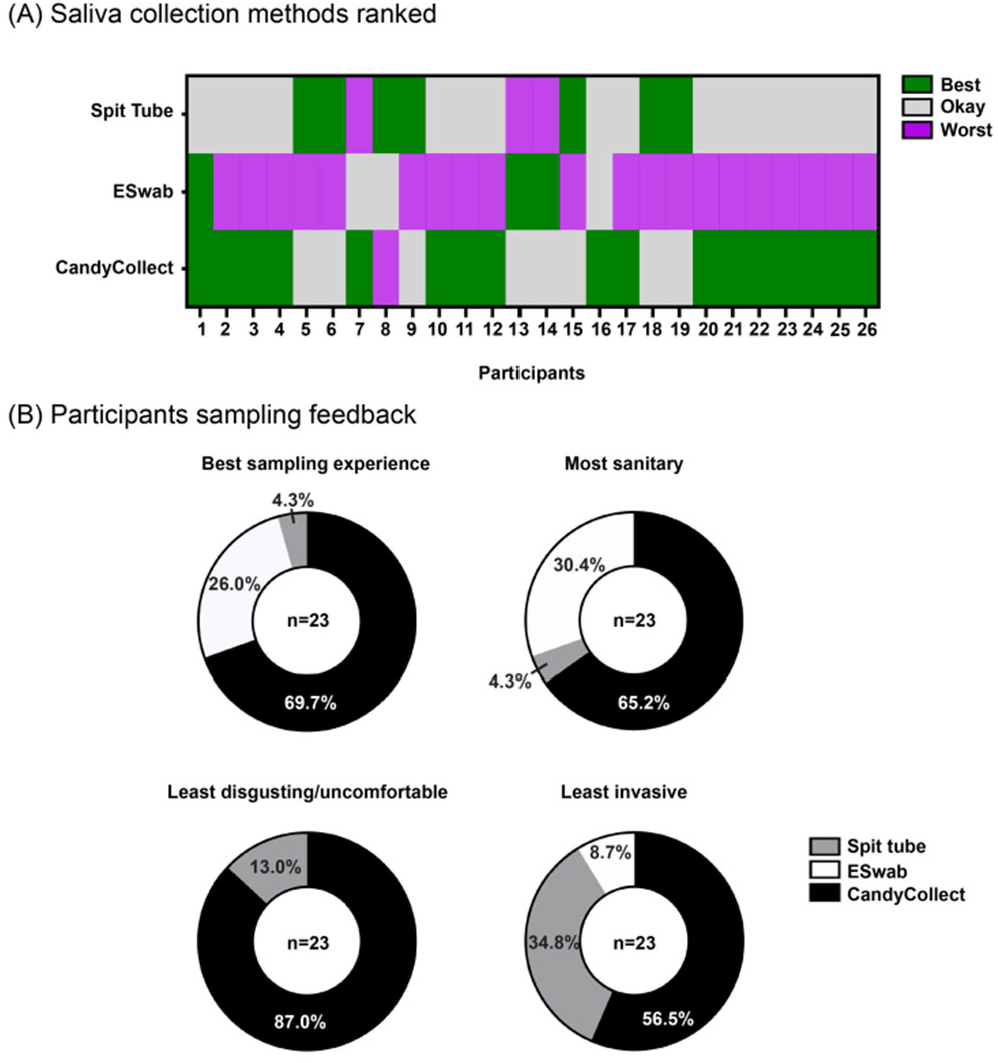
Participant feedback shows overall preference for CandyCollect devices. (A) Participants were asked to rank the three sampling methods (CandyCollect, ESwab™, and Spit Tube) in terms of best overall sampling method to worst overall sampling method. (B) Participants were asked to select one sampling method that most accurately fits the above descriptions (best sampling experience, most sanitary, least disgusting/uncomfortable, and least invasive). The CandyCollect device was the most frequently selected sampling method for all of the above user feedback questions.

### Human subject sample processing

Participants stored their samples at ambient temperature, and they were picked up the following day using United Parcel Service (UPS) Next Day Air. Samples were stored at -20 °C upon receipt and transferred to -80 °C for longer term storage before processing. All laboratory procedures were performed in accordance with Biosafety Level-2 laboratory practices and the University of Washington Site-Specific Bloodborne Pathogen Exposure Control Plan. *S. mutans* and *S. aureus* on CandyCollect devices were eluted and lysed following the protocol stated above. To avoid unnecessary freeze-thaw cycles, ESwab™ and SpeciMAX Stabilized Saliva Collection Kits samples were aliquoted into 20 μL aliquots and stored at -80 °C. For ESwab™ and SpeciMAX Stabilized Saliva Collection Kits samples, DNA was isolated using MagMAX™ Total Nucleic Acid Isolation Kit (ThermoFisher Scientific, Cat# AM1840) according to the protocol “Disruption of liquid samples” supplied by the manufacturer. Briefly, 175 μL of aliquoted samples were transferred to each bead beating tube provided in the kit followed by the addition of 230 μL of Lysis/Binding solution. Bead beating was carried out via MiniBeadBeater (mentioned above) twice for 30 s, then each tube was centrifuged at 16,000 g for 3 min. Afterward, genomic DNA of *S. mutans* and *S. aureus* was isolated and enriched following the protocol stated above and quantified using qPCR with a detection limit of 25 fg.

### Human subject data analysis

Conversion factor: DNA content reported by the qPCR analysis was converted to estimated copy number of bacterial DNA/mL to facilitate comparisons between the three methods of saliva collection: CandyCollect, ESwab™, and SpeciMAX Stabilized Saliva Collection Kits. A conversion factor to derive bacterial concentrations/DNA content was calculated for each method, taking into consideration the variations in the collection and processing.

In brief, qPCR results from the CFX connect Real-Time PCR Detection System were reported in ng of DNA corresponding to the 5 μL samples loaded into the machine. Dilutions corresponding to the purification and enrichment of samples (see Methods section “Isolation, purification, and enrichment of genomic DNA from *S. mutans, S. aureus*, and *S. pyogenes*”) were used to derive the DNA content of samples input into the purification and enrichment procedure. Estimates for sample dilution during sampling were employed to further convert the qPCR output reading to ng DNA per mL saliva. These sample dilution estimates are as follows: Spit tube, we assumed participants provided 1 mL of saliva as instructed into a tube containing 1 mL of stabilization solution; ESwab™, we assumed 130 μL of saliva is captured during sampling which was then stored in 1 mL of ESwab™ buffer; CandyCollect, we assumed 50 μL of saliva was captured by device. To report copy number of bacterial DNA/mL, based on the genome sizes of the bacteria (2.6 fg / genome for *S. mutans*, 3.0 fg /genome for *S. aureus*),^26, 27^ detected DNA values (ng) were converted to the equivalent bacteria number and the data were reported estimated copy number of bacterial DNA/mL.

Statistics: Statistical analysis was performed using GraphPad Prism 9 software. One-way analysis of variance (One-way ANOVA) was chosen to compare groups and Tukey’s multiple comparison tests were further used in evaluating significance of pairwise comparisons.

### Shelf life test

The shelf life experiment followed the protocol established in our previous work.^1^ In brief, the CandyCollect devices were plasma treated (see Methods section “CandyCollect device fabrication”) in descending order 1 year (369 days), 4 months (137 days), 3 months (105 days) and 0 days (control) prior to the experiment with n = 3 replicated devices per time point and concentration. This allowed for all devices to be tested on the same day. Each time point was tested in triplicate at four concentrations of *S. pyogenes*, 10^3^, 10^4^, 10^5^ and 10^9^ CFU/mL, in addition to negative controls (concentration 0 CFU/mL).

## Results and Discussion

### Capture, Elution, and qPCR detection of *S. mutans* and *S. aureus* from CandyCollect Devices

In testing the capture ability of CandyCollect device with bacteria other than *S. pyogenes*, which was reported in our previous work,^1^ finding an elution method that worked for all bacteria of interest, *S. pyogenes, S. mutans*, and *S. aureus*, was important to establish the versatility of the device. Effective elution is crucial for accurate analysis by qPCR. Various elution buffers were accessed for high elution rates. Fluorescence images were first quantified to assess the elution efficiency of the elution buffers (Figure 3A and S6), with high elution efficiency resulting in less fluorescent signal after elution. While some elution buffers demonstrated low elution efficiency (ESwab™ buffer with 5% ethanol, PBS with 2% SDS, and ESwab™ buffer with 2% SDS), those that demonstrated high elution efficiency (PBS with 5% Proteinase K and ESwab™ buffer with 5% Proteinase K) were then evaluated through qPCR. Several elution buffers were further eliminated when qPCR results demonstrated some of elution buffers contained qPCR inhibitors (data not shown), thus preventing downstream analysis of CandyCollect samples. We selected PBS with 5% Proteinase K as the elution buffer for the CandyCollect device as it was the buffer with the highest observed elution rates: approximately 90% of *S. mutans* and 95% of *S. aureus* were removed from CandyCollect (Figure 3Ai and 3Aii).

We also established the qPCR assays for analyzing DNA content from eluted *S. mutans* and *S. aureus* samples (see Methods section “Quantitative PCR assay for detection of *S. mutans, S. aureus*, and *S. pyogenes*” and Supplementary Information), which yielded good linear relationships between the DNA content from both eluted bacteria samples and bacterial concentrations incubated on the device (Figure 3B). The elution buffer and qPCR assay were also tested on a solution containing a mixture of *S. mutans, S. aureus* and *S. pyogenes* from CandyCollect devices (Figure S7). The result showed that PBS with 5% Proteinase K was able to elute the samples containing multiple bacteria (Figure S7).

### Analysis of Human Subjects Samples via qPCR Demonstrates Feasibility of CandyCollect Devices for Salivary Commensal Bacteria Capture

We compared the CandyCollect device to two commercially available methods for oral sample/saliva collection, ESwab™ (oral swab) and SpeciMAX Stabilized Saliva Collection Kits (spit tube). Participants were instructed to provide a sample using the three methods as follows: CandyCollect (suck the samples until candy/flavor is gone), ESwab™ (suck 30 seconds), and Spit tube (collect 1 mL saliva). Although capture of bacterial pathogens is the ultimate goal for the CandyCollect device, commensal bacteria—bacteria present in healthy hosts—were selected as a proof-of-concept analytes to evaluate the device in order to maximize the population available for enrollment in this initial study. Individuals have different populations of commensal bacteria in their oral microbiome. As such, *S. mutans* and *S. aureus* are not universally present in the population, so we did not expect to detect these bacteria in samples from all participants. In general, *S. mutans* is more prevalent in the population compared to *S. aureus*, at 80-87% and 18-39% prevalence, respectively.^16-19^ As such, it is unsurprising that more participants had detectable *S. mutans* than *S. aureus*.

Importantly, for participants in which a given bacterium (*S. mutans* or *S. aureus*) was detected in one or both of the commercially available methods (ESwab™ and Spit tubes), CandyCollect devices had a 100% concordance with the other positive results (Figure 4). As expected in human subjects studies, there was some variability in positive/negative results across sampling methods and across sampling days. For example, *S. aureus* was detected in Participant 8 on both Day 1 and 2 in the ESwab™ and CandyCollect samples, but not in Spit tubes sample. In Participant 4, on Day 2, *S. aureus* was detected in the spit tube and the CandyCollect samples, but not in the ESwab™ sample. While we cannot identify the exact cause of this variability, it is noteworthy that the CandyCollect device did not fail to capture *S. mutans* or *S. aureus* when they were collected by either of the two commercially available methods. In Figure 4, we have reported the data as estimated copy number of bacterial DNA/mL in the original saliva sample; this concentration is an estimate based on an estimated volume of saliva collected by the CandyCollect device and ESwab™ (as described in the Methods section). However, due to approximations in the collection volume these concentrations are less relevant than the presence or absence of the bacteria.

### User Feedback Indicates CandyCollect Devices are the Preferred Saliva Sampling Tool

Of the 26 participants that completed the electronic survey, 65% of them selected CandyCollect as the best method of saliva collection (Figure 5A). Of the 23 participants who completed the detailed survey questions (Figure 5B), 70% of the participants ranked CandyCollect as being the best sampling experience, 65% chose it as the most sanitary sampling method, 87% selected it as being the least disgusting and uncomfortable, and 57% selected it as being the least invasive. (Note on sample size in Figure 5: Figure 5A represents n=26 and Figure 5B represents n=23 due to an electronic survey error wherein 3 participants failed to receive all survey questions (see Methods section “Human subjects’ sample and feedback collection”). Overall, the CandyCollect device was selected by the majority of participants as the preferred saliva collection method (Figure 5A) and sampling experience (Figure 5B).

### Shelf Life Tests: CandyCollect Devices Capture *S. pyogenes* after 1 Year of Storage

Bacterial adhesion to the polystyrene channel of the CandyCollect device is facilitated by an increase in the hydrophilicity and wettability caused by oxygen plasma treatment of the surface.^1,28-29^ Hydrophilicity has been observed to decay over time,^30^ potentially reducing the efficacy of bacterial capture by the CandyCollect device. Previously, we established through quantification of fluorescence images, that there was no notable difference in bacterial adhesion to CandyCollect devices plasma treated 0, 3, 7 and 14 days before bacterial incubation, and devices plasma treated 0 and 62 days before bacterial incubation.^1^ Here, we have expanded on this research by evaluating CandyCollect devices over longer time frames, which is required for the device to be effective in a commercial setting. Devices from all time points—0 days, 3 months, 4 months and 1 year—were able to collect bacteria (Figure 6, S8). Although there is a significant difference between the integrated density per area in the images from the 0 days and 1-year devices (Figure 6B), there is only a 25% reduction in bacteria captured after 1 year of storage. In the future, longer lasting surface treatment can be used, such as treatments used for commercially available cell cultureware, which typically has a multiyear shelf life.

**Figure 6.**
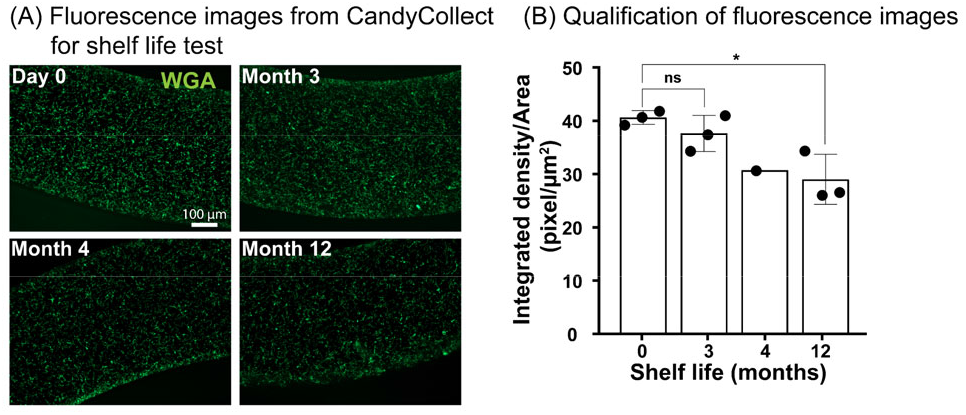
Shelf life tests demonstrate that CandyCollect devices effectively capture *S. pyogenes* after 1 year of storage. Devices were plasma treated and stored at room temperature for 0 days (control group), 3 months, 4 months, and 1 year. (A) Fluorescence microscopy images indicate capture of S. pyogenes after 1 year of storage is similar to the control, with ∼25% decrease in bacteria captured. (B) Quantification of the integrated density per area (pixel/μm^2^). Data sets were analyzed using one-way ANOVA and Tukey’s multiple comparison tests (*p<0.05). No significant difference between 0 days (control group) and 3 months of storage was found. Note: depicted images are from CandyCollect devices incubated with *S. pyogenes* at a concentration of 1×10^9^ CFU/mL for 10 minutes. Each data point represents an individual CandyCollect device (4 images were taken per device, and the data point plotted is the average); the bars represent the mean ± SEM of n=3 CandyCollects. *S. pyogenes* was fluorescently labeled with WGA.

While *S. mutans* and *S. aureus* are well established commensal microbes, it is important to note that the location of these bacteria within the oropharyngeal space might differ from the location of pathogens of interest. GAS-related strep throat is an infection of the back of the throat. *S. mutans* is a commensal bacterium of the oral cavity and particularly the gingiva (i.e., gums).^31, 32^ *S. aureus* is located primarily in the anterior nares and is also found in other places in the respiratory tract (e.g., mouth, nose, and throat).^33^ Nevertheless, independent of the primary location of each microbe, it is known that these bacteria are present in saliva.^34, 35^ The present study lays the foundation for microbial collection from human subjects using the CandyCollect device, and it is important to evaluate the device in subsequent clinical studies for each pathogen of interest.

We acknowledge that this pilot study was performed with a relatively small sample size; the goal was to establish the potential of the CandyCollect device in healthy adults before progressing to individuals with respiratory illness. We have ongoing studies with larger sample sizes in adults and children with respiratory illness. These additional studies also include user feedback surveys, and we will determine if the feedback from these larger cohorts is consistent with the feedback from this initial cohort.

### Conclusion

In this work, we used two commensal bacteria, *S. mutans* and *S. aureus*, as proof-of-concept bacteria to demonstrate the abilities of the CandyCollect device in capturing salivary bacteria in healthy adults. The results showed that (1) the CandyCollect device can effectively capture commensal bacteria from healthy participants in a home setting, (2) samples are stable through standard shipping at room temperature and the bacteria can be eluted and quantified using qPCR, (3) most users ranked CandyCollect as their first choice for oral sam-pling method (compared to standard oral swabs and spit tubes), (4) the CandyCollect device is functional after storage times of up to one year. For more reproducible clinical characterization work and commercial implementation, simple manufacturing can be set up by using rapid injection molding as the design is fully amenable to injection molding.^36^ The present study opens up several exciting areas of future work as a new tool for at-home and in-clinic sampling that is intuitive, convenient, and child friendly. Currently, we are conducting a study using CandyCollect with patients age 5-17 with GAS pharyngitis. In addition, future work includes extending the capabilities of CandyCollect to viruses, mycobacteria, and fungal pathogens.

## ASSOCIATED CONTENT

### Supporting Information

The supporting information for publication includes additional data on the modification and verification of primers for *S. aureus* qPCR analysis, standard curves for human subject sample qPCR assays, experiments testing various elution buffers, qPCR tests validating the method on a mixture of three bacteria in saliva, and experiments to evaluate the shelf life of the CandyCollect device.

## AUTHOR INFORMATION

### Authors

**Wan-chen Tu** − *Department of Chemistry, University of Washington, Seattle, Washington 98195, United States*

**Anika M. McManamen** − *Department of Chemistry, University of Washington, Seattle, Washington 98195, United States*

**Xiaojing Su** − *Department of Chemistry, University of Washington, Seattle, Washington 98195, United States*

**Ingrid Jeacopello** − *Department of Chemistry, University of Washington, Seattle, Washington 98195, United States*

**Meg G. Takezawa** − *Department of Chemistry, University of Washington, Seattle, Washington 98195, United States*

**Damielle L. Hieber** − *Department of Chemistry, University of Washington, Seattle, Washington 98195, United States*

**Grant W. Hassan** − *Department of Chemistry, University of Washington, Seattle, Washington 98195, United States*

**Ulri N. Lee** − *Department of Chemistry, University of Washington, Seattle, Washington 98195, United States*

**Eden V. Anana** − *Department of Chemistry, University of Washington, Seattle, Washington 98195, United States*

**Mason P. Locknane** − *Department of Chemistry, University of Washington, Seattle, Washington 98195, United States*

**Molly W. Stephen-son** − *Department of Chemistry, University of Washington, Seattle, Washington 98195, United States*

**Victoria A. M. Shinkawa** − *Department of Chemistry, University of Washington, Seattle, Washington 98195, United States*

**Ellen R. Wald** − *Department of Pediatrics, University of Wisconsin School of Medicine and Public Health, Madison, Wisconsin 53792, USA*

**Gregory P. DeMuri** − *Department of Pediatrics, University of Wisconsin School of Medicine and Public Health, Madison, Wisconsin 53792, USA*

**Karen Adams** − *Institute of Translational Health Sciences, School of Medicine, University of Washington, Seattle, Washington 98109, USA*

**Erwin Berthier** − *Department of Chemistry, University of Washington, Seattle, Washington 98195, United States*

## Author Contributions

WCT, AMM, XS, ERW, GPD, KNA, EB, ST, and ABT conceptualized the research. UNL designed CandyCollect devices. AMM, EVA, and WCT milled CandyCollect devices. DLH, EVA and MWS fabricated candy devices and prepared devices for human subject research. WCT, AMM, XS, MGT, and GWH reviewed the literature that informed the study. WCT, AMM, and XS designed the biological experiments. WCT, AMM, XS, and IJ conducted biological experiments and data collection. KNA advised on work with human subjects study design and regulatory protocols. IJ, MGT, GWH, MPL, and VAMS recruited participants, set up the REDCap platform to screen participant eligibility and administer survey questions, packaged and shipped CandyCollects to research participants, and communicated with participants. WCT, AMM, XS, EB, ST, and ABT interpreted the data. WCT, AMM, XS, MPL, ST, and ABT wrote sections of the manuscript. WCT, AMM, XS, and MPL made figures for the manuscript. ERW and GPD provided expertise on clinical relevance and sampling bacteria in saliva. WCT, AMM, XS, MPL, ERW, GPD, KNA, EB, ST, and ABT edited and revised the manuscript, and all authors approved the manuscript. ABT and ST supervised the research.

## ACKNOWLEDGMENT

This work was supported by National Institutes of Health grants (R21AI166120, R35GM128648 (the latter specifically supported some of the in-lab developments and in vitro experiments)), The Camille and Henry Dreyfus Foundation, an Alfred P. Sloan Research Fellowship, the David and Lucile Packard Foundation, the Society for Laboratory Automation and Screening (SLASFG2020, UNL), Ronald E. McNair Scholars (DLH), Louis Stokes Alliance for Minority Participation (DLH), and the University of Washington. REDCap at UW ITHS is supported by the National Center For Advancing Translational Sciences of the National Institutes of Health under Award Number UL1 TR002319. The content is solely the responsibility of the authors and does not necessarily represent the official views of the Society for Laboratory Automation and Screening, the National Institutes of Health, or other funding sources.

We would also like to thank Lochlan Hickok and Paul Miller for helping facilitate shipping and logistics; Yixuan Zhou, Aryam Chhazal, Ben Mous, and Juan C. Sanchez for helping with device fabrication; and David N. Phan for making the Schematic of the CandyCollect device dimensions. Moreover, we thank the participants enrolled in this study.

## CONFLICTS OF INTEREST

ABT has ownership in Stacks to the Future, LLC; ST: ownership in Stacks to the Future, LLC, Tasso, Inc., and Salus Discovery, LLC; and EB has ownership in Stacks to the Future, LLC, Tasso, Inc., and Salus Discovery, LLC and employment by Tasso, Inc. However, this research is not related to these companies.

## Notes

### Summary of Updates

We added one more figure in the Supplementl material and also added one more funding which we forgot previously.

## REFERENCES

(1) Lee, U. N.; Su, X.; Hieber, D. L.; Tu, W. C.; McManamen, A. M.; Takezawa, M. G.; Hassan, G. W.; Chan, T. C.; Adams, K. N.; Wald, E. R.; DeMuri, G. P.; Berthier, E.; Theberge, A. B.; Thongpang, S. CandyCollect: at-home saliva sampling for capture of respiratory pathogens. Lab Chip 2022, 22 (18), 3555–3564.

(2) Zar H. J.; Ferkol, T. W. The global burden of respiratory disease—Impact on child health. Pediatr. Pulmonol. 2014, 49, 430–434.

(3) Bisno, A. L. Acute Pharyngitis. N. Engl. J. Med. 2001, 344, 205–211

(4) Kaitz .; Sabato, R.; Shalev, I.; Ebstein, R.; Mankuta, D. Children’s noncompliance during saliva collection predicts measures of salivary cortisol. Dev. Psychobiol. 2012, 54, 113–123.

(5) Shulman, T.; Bisno, A. L.; Clegg, H. W.; Gerber, M. A.; Kaplan, E. L.; Lee, G.; Martin, J. M.; Van Beneden, C. Clinical Practice Guideline for the Diagnosis and Management of Group A Streptococcal Pharyngitis: 2012 Update by the Infectious Diseases Society of America. Clin. Infect. Dis. 2012, 55, e86–e102.

(6) Piasio, N.; Turner, N.; Wheeler, A. Methods and devices for testing saliva. WIPO (PCT) 2010/0273.177 A1, 2007.

(7) DeMuri, G.; Wald, E. R. Detection of Group A Streptococcus in the Saliva of Children Presenting With Pharyngitis Using the cobas Liat PCR System. Clin. Pediatr. 2020, 59, 856–858.

(8) Comber, L.; Walsh, K. A.; Jordan, K.; O’Brien, K. K.; Clyne, B.; Teljeur, C.; Drummond, L.; Carty, P. G.; De Gascun, C. F.; Smith, S. M.; Harrington, P.; Ryan, M.; O’Neill, M. Alternative clinical specimens for the detection of SARS-CoV-2: A rapid review. Rev. Med. Virol. 2021, 31, e2185.

(9) Heat inactivation of respiratory viruses in raw saliva for nucleic acid extraction. ThermoFisher.https://assets.thermofisher.com/TFS-Assets/BID/Application-Notes/heat-inactivation-viruses-nucleic-acid-extraction-app-note.pdf (accessed 2022/12/29).

(10) Saliva Collection and Handling Advice. 3 rd Edition. Salimetrics. https://www.researchgate.net/profile/Nirmala-Svsg/post/Saliva-collection-in-rats/attachment/59d633c579197b807799173e/AS%3A376127236395008%401466687129091/download/Saliva_Collection_Handbook.pdf (accessed 2022/12/29).

(11) Melo Costa, M.; Benoit, N.; Dormoi, J.; Amalvict, R.; Gomez, N.; Tissot-Dupont, H.; Million, M.; Pradines, B.; Granjeaud, S.; Almeras, L. Salivette, a relevant saliva sampling device for SARS-CoV-2 detection. J. Oral Microbiol. 2021, 13, 1920226.

(12) Ottaviano E.; Parodi C.; Borghi E.; Massa V.; Gervasini C.; Centanni S.; Zuccotti G.; Bianchi S. Saliva detection of SARS-CoV-2 for mitigating company outbreaks: a surveillance experience, Milan, Italy, March 2021. Epidemiol. Infect. 2021, 149: e171.

(13) De Meyer J.; Goris H.; Mortelé O.; Spiessens A.; Hans G.; Jansens H.; Goossens H.; Matheeussen V.; Vandamme S. Evaluation of Saliva as a Matrix for RT-PCR Analysis and Two Rapid Antigen Tests for the Detection of SARS-CoV-2. Viruses. 2022; 14(9):1931.

(14) Martín, R.; Miquel, S.; Ulmer, J.; Kechaou, N.; Langella, P.; Bermúdez-Humarán, L. G. Role of commensal and probiotic bacteria in human health: a focus on inflammatory bowel disease. Microb. Cell Fact. 2013, 12, 71.

(15) Kreth, J.; Giacaman, R. A.; Raghavan, R.; Merritt, J. The road less traveled – defining molecular commensalism with Streptococcus sanguinis. Mol. Oral Microbiol. 2017, 32, 181–196.

(16) McCormack, M. G.; Smith, A. J.; Akram, A. N.; Jackson, M.; Robertson, D.; Edwards, G. Staphylococcus aureus and the oral cavity: An overlooked source of carriage and infection? Am. J. Infect. Control 2015, 43 (1), 35–37.

(17) Ohara-Nemoto, Y.; Haraga, H.; Kimura, S.; Nemoto, T. K. Occurrence of staphylococci in the oral cavities of healthy adults and nasal–oral trafficking of the bacteria. J. Med. Microbiol. 2008, 57, 95–99

(18) Pannu, P.; Gambhir, R.; Sujlana, A. Correlation between the salivary Streptococcus mutans levels and dental caries experience in adult population of Chandigarh, India. Eur J Dent 2013, 07, 191–195.

(19) Lee Y. J; Kim, M.-A.; Kim, J.-G.; Kim, J.-H. Detection of Streptococcus mutans in human saliva and plaque using selective media, polymerase chain reaction, and monoclonal antibodies. Oral Biol Res 2019, 43, 121–129.

(20) Koblitz, J.; Halama, P.; Spring, S.; Thiel, V.; Baschien, C.; Hahnke, Richard L.; Pester, M.; Overmann, J.; Reimer, Lorenz C. MediaDive: the expert-curated cultivation media database. Nucleic Acids Res. 2022, 51 (D1), D1531–D1538.

(21) ATCC Medium: 18 Tryptic Soy Agar/Broth (Soybean-Casein Digest Medium, USP) https://www.atcc.org/-/media/product-assets/documents/microbial-media-formulations/1/8/atcc-medium-18.pdf?rev=832846e1425841f19fc70569848edae7 (accessed 2022/8/20).

(22) Gera, K. and McIver, K. S. Laboratory Growth and Maintenance of Streptococcus pyogenes (The Group A Streptococcus, GAS). Curr. Protoc. Microbiol. 2013, 30, 9D.2.1–9D.2.13.

(23) Lochman, J.; Zapletalova, M.; Poskerova, H.; Izakovicova Holla, L.; Borilova Linhartova, P. Rapid Multiplex Real-Time PCR Method for the Detection and Quantification of Selected Cariogenic and Periodontal Bacteria. Diagnostics 2020, 10, 8

(24) Wood, C.; Sahl, J.; Maltinsky, S.; Coyne, B.; Russakoff, B.; Yagüe, D. P.; Bowers, J.; Pearson, T. SaQuant: a real-time PCR assay for quantitative assessment of Staphylococcus aureus. BMC Microbiol. 2021, 21, 174.

(25) Galia, L.; Ligozzi, M.; Bertoncelli, A.; Mazzariol, A. Real-time PCR assay for detection of Staphylococcus aureus, Panton-Valentine Leucocidin and Methicillin Resistance directly from clinical samples. AIMS Microbiol. 2019, 5, 138–146.

(26) Yano A.; Kaneko N.; Ida H.; Yamaguchi T.; Hanada N. Real-time PCR for quantification of Streptococcus mutans. FEMS Microbiol Lett. 2002 19, 217, 23–30.

(27) Yang S.; Lin S.; Kelen G. D.; Quinn T. C.; Dick J. D.; Gaydos C. A.; Rothman R. E. Quantitative multiprobe PCR assay for simultaneous detection and identification to species level of bacterial pathogens. J Clin Microbiol. 2002, 40, 3449–54

(28) Hemmatian, T.; Lee, H.; Kim, J. Bacteria Adhesion of Textiles Influenced by Wettability and Pore Characteristics of Fibrous Substrates. Polymers 2021, 13, 223.

(29) Guruvenket, S.; Rao, G. M.; Komath, M.; Raichur, A. M. Plasma surface modification of polystyrene and polyethylene. Appl. Surf. Sci. 2004, 236, 278–284.

(30) Li, J.; Oh, K.; Yu, H. Surface rearrangements of oxygen plasma treated polystyrene: surface dynamics and humidity effect. Chin. J. Polym. Sci. 2005, 23, 187–196.

(31) Duchin, S.; Van Houte, J. Relationship of Streptococcus mutans and lactobacilli to incipient smooth surface dental caries in man. Arch. Oral Biol. 1978, 23 (9), 779–786.

(32) Berlutti, F.; Catizone, A.; Ricci, G.; Frioni, A.; Natalizi, T.; Valenti, P.; Polimeni, A. Streptococcus mutans and Streptococcus sobrinus are able to adhere and invade human gingival fibroblast cell line. Int. J. Immunopathol. Pharmacol. 2010, 23 (4), 1253–1260.

(33) Nilsson, P.; Ripa, T. Staphylococcus aureus throat colonization is more frequent than colonization in the anterior nares. J. Clin. Microbiol. 2006, 44, 9, 3334–3339.

(34) Köhler, B.; Bratthall, D. Practical method to facilitate estimation of Streptococcus mutans levels in saliva. J. Clin. Microbiol. 1979, 9 (5), 584–588.

(35) Petti, S.; Boss, M.; Messano, G. A.; Protano, C.; Polimeni, A. High salivary Staphylococcus aureus carriage rate among healthy paedodontic patients. New Microbiol. 2014, 37 (1), 91–96.

(36) Lee, U. N.; Su, X.; Guckenberger, D. J.; Dostie, A. M.; Zhang, T.; Berthier, E.; Theberge, A. B. Fundamentals of rapid injection molding for microfluidic cell-based assays. Lab Chip 2018, 18, 496–504.

