## supplemental material for "At-home saliva sampling in healthy adults using CandyCollect, a lollipop-inspired device"

**Table S1.** Mass, diameter, thickness, and dissolving time of CandyCollects in Figure 4 (Day 1)

**Table S2.** Mass, diameter, thickness, and dissolving time of CandyCollects in Figure 4 (Day 2)

**Table S3.** The sequences of *S. aureus* primers and probe

**Table S4.** Survey questions for user feedback

**Figure S1**: Schematic of the CandyCollect device dimensions

**Figure S2**: *S. aureus* primer validation by qPCR with SYBR Green chemistry

**Figure S3**: *S. aureus* primer validation by agarose gel electrophoresis and qPCR with TaqMan probe

**Figure S4:** Standard curves for the *S. mutans* and *S. aureus* qPCR assay

**Figure S5:** At-home saliva sampling devices for human subjects study

**Figure S6:** Additional elution experiments demonstrate that *S. aureus* captured by CandyCollect devices can be removed efficiently via elution buffers

**Figure S7:** qPCR tests demonstrate that the mixture of three bacteria in saliva can be analyzed for their individual concentrations

**Figure S8:** qPCR shelf life tests demonstrate that the CandyCollect device effectively captures *S. pyogenes* after 1 year of storage

**Table S1.** Mass, diameter, thickness, and dissolving time of CandyCollects in Figure 4 (Day 1)

| Participant | Mass (g) | Diameter (mm) | Thickness (mm) | Time (min) |
|---|---|---|---|---|
| 1 | 1.34 | 16 | 4 | 4:35 |
| | 1.24 | 16 | 4 | 4:16 |
| | 1.37 | 16 | 4 | 5:00 |
| 2 | 1.26 | 16 | 4 | 2:48 |
| | 1.21 | 16 | 4 | 3:31 |
| | 1.38 | 16 | 4 | 3:51 |
| 3 | 1.34 | 16 | 4 | 4:13 |
| | 1.33 | 16 | 4 | 5:01 |
| | 1.28 | 16 | 4 | 4:16 |
| 4 | 1.37 | 16 | 4 | 4:50 |
| | 1.23 | 16 | 4 | 4:21 |
| | 1.40 | 16 | 4 | 4:34 |
| 5 | 1.34 | 16 | 4 | 4:25 |
| | 1.30 | 16 | 4 | 4:51 |
| | 1.40 | 16 | 4 | 4:31 |
| 6 | 1.32 | 16 | 4 | 5:02 |
| | 1.39 | 16 | 4 | 5:37 |
| | 1.31 | 16 | 4 | 4:19 |
| 7 | 1.24 | 16 | 4 | 8:32 |
| | 1.41 | 16 | 4 | 7:00 |
| | 1.32 | 16 | 4 | 7:52 |
| 8 | 1.30 | 16 | 4 | 7:58 |
| | 1.39 | 16 | 4 | 8:22 |
| | 1.34 | 16 | 4 | 4:43 |

| | | | | |
|---|---|---|---|---|
| 9 | 1.40 | 16 | 4 | 2:48 |
| | 1.23 | 16 | 4 | 2:10 |
| | 1.36 | 16 | 4 | 2:12 |
| 10 | 1.86 | 16 | 5 | 3:14 |
| | 1.86 | 16 | 5 | 2:34 |
| | 1.90 | 16 | 5 | 2:45 |
| 11 | 1.33 | 16 | 4 | 5:04 |
| | 1.37 | 16 | 4 | 3:58 |
| | 1.37 | 16 | 4 | 4:54 |
| 12 | 1.40 | 16 | 4 | 2:37 |
| | 1.34 | 16 | 4 | 2:06 |
| | 1.21 | 16 | 4 | 2:39 |
| 13 | 1.24 | 16 | 4 | 4:35 |
| | 1.33 | 16 | 4 | 4:32 |
| | 1.21 | 16 | 4 | 4:25 |
| 14 | 1.32 | 16 | 4 | 5:41 |
| | 1.32 | 16 | 4 | 5:24 |
| | 1.32 | 16 | 4 | 5:31 |

**Table S2.** Mass, diameter, thickness, and dissolving time of CandyCollects in Figure 4 (Day 2)

| **Participant** | **Mass (g)** | **Diameter (mm)** | **Thickness (mm)** | **Time (min)** |
|---|---|---|---|---|
| 1 | 1.7 | 16 | 5 | 4:45 |
| | 1.72 | 16 | 5 | 4:33 |
| | 1.83 | 16 | 5 | 5:00 |
| 2 | 1.80 | 16 | 5 | 3:51 |
| | 1.76 | 16 | 5 | 2:49 |
| | 1.77 | 16 | 5 | 3:29 |
| 3 | 1.70 | 16 | 5 | 5:46 |
| | 1.72 | 16 | 5 | 6:20 |
| | 1.72 | 16 | 5 | 5:54 |
| 4 | 1.84 | 16 | 5 | 5:50 |
| | 1.79 | 16 | 5 | 5:00 |
| | 1.76 | 16 | 5 | 4:59 |
| 5 | 1.77 | 16 | 5 | 6:01 |
| | 1.75 | 16 | 5 | 5:20 |
| | 1.72 | 16 | 5 | 4:48 |
| 6 | 1.82 | 16 | 5 | 4:53 |
| | 1.88 | 16 | 5 | 5:01 |
| | 1.9 | 16 | 5 | 4:21 |
| 7 | 1.81 | 16 | 5 | 9:08 |
| | 1.83 | 16 | 5 | 10:34 |
| | 1.77 | 16 | 5 | 9:49 |
| 8 | 1.74 | 16 | 5 | 4:53 |
| | 1.81 | 16 | 5 | 6:18 |
| | 1.90 | 16 | 5 | 4:18 |

| | | | | |
|---|---|---|---|---|
| 9 | 1.86 | 16 | 5 | 2:47 |
| | 1.81 | 16 | 5 | 2:34 |
| | 1.89 | 16 | 5 | 2:39 |
| 10 | 1.31 | 16 | 4 | 2:08 |
| | 1.24 | 16 | 4 | 2:57 |
| | 1.37 | 16 | 4 | 3:10 |
| 11 | 1.89 | 16 | 5 | 5:35 |
| | 1.77 | 16 | 5 | 6:02 |
| | 1.76 | 16 | 5 | 6:49 |
| 12 | 1.83 | 16 | 5 | 4:37 |
| | 1.81 | 16 | 5 | 4:49 |
| | 1.77 | 16 | 5 | 4:59 |
| 13 | 1.86 | 16 | 5 | 5:37 |
| | 1.77 | 16 | 5 | 6:11 |
| | 1.88 | 16 | 5 | 6:35 |
| 14 | 1.81 | 16 | 5 | 5:32 |
| | 1.9 | 16 | 5 | 6:22 |
| | 1.79 | 16 | 5 | 6:36 |

***S. aureus* primer modification and verification**

The primer/probe sequences for the *S. aureus* qPCR assay were referenced from Galia et al., 2019[1] with minor modifications for both forward and reverse primers. Probe sequence remains the same as in Galia et al., 2019[1] except using FAM as a reporter dye. The forward primer sequence was modified based on the NCBI database for *S. aureus* sequence (25923) (GenBank accession no. CP000046); the reverse primer sequence modification was based on the ATCC Genomes database for *S. aureus* (25923). All primer/probe sequences are listed in Table S1 below. Primer validation is shown in Figure S1.

**Table S3.** The sequences of S. aureus primers and probe. The original forward (F0), reverse (R0) primers, and probe are adopted from Galia et al., 2019; in this study, modified forward and reverse primers were designated as F1 and R1.

| **Primers/probe** | **sequences** | **Notes** |
|---|---|---|
| Forward (F0) | 5'-GGCATATGTATGGCAATTGTTTC-3' | Galia et al. 2019[1] |
| Forward (F1) | 5'-GGCATATGTATGGCAATCGTTTC-3' | This study |
| Reverse (R0) | 5'-CGTATTGCCCTTTCGAAACATT-3' | Galia et al. 2019[1] |
| Reverse (R1) | 5'-CGTATTGTTCTTTCGAAACATT-3' | This study |
| Probe | 5'-/56-FAM/ATT ACT TAT AGG GAT GGC TAT C/3MGB-NFQ/-3' | Galia et al. 2019[1] |

We first tested the original forward/reverse primer pair (F0/R0), from Galia et al., 2019,[1] for qPCR amplification and efficiency with SYBR Green chemistry. A 1:10 serial dilution of purified DNA from *S. aureus* was used as templates (10 ng to 10 fg/reaction). Purified DNA from *S. mutans* and *S. pyogenes* (10 ng/reaction) were used as negative controls. qPCR was run under the following thermal cycling conditions: 5 min at 95 °C followed by 40 cycles of 95 °C for 15 s and 60 °C for 30 s. Based on the amplification plot (Figure S1), qPCR with F0/R0 primer pair did not have a good dynamic range of detection and the reaction efficiency was low. In addition, two negative controls had strong fluorescence signals (data not shown). This prompted us to seek improvement on primer design. The modified forward primer sequence (F1) has only one nucleotide different from the F0; the modified reverse primer sequence (R1) has two nucleotides different from the R0 (Table S1). To validate the new primers, the new primer was paired with the original forward or reverse primers or another new primer: F0/R1, F1/R0, F1/R1, and was examined for qPCR amplification and efficiency as above. The F0/R1 primer pair had improved assay dynamic range and efficiency compared to the original F0/R0. The F1/R0 and F1/R1 primer pairs sets provided the best assay dynamic range and efficiency (Figure S1). It is noted that non-specific amplification can also be detected in the qPCR assays for these three pairs of primers. To increase specificity, F1/R0 and F1/R1 primer pairs were combined with the probe originally designed from Galia et al., 2019[1], and TaqMan master mix was used for the qPCR assay with purified DNA as above. The results showed that the F1/R1 primer pair had better sensitivity and wider assay dynamic range compared with F1/R0 pair (Figure S2A). The efficiency is 85%. Agarose gel electrophoresis showed a single PCR product (~70 bp, an expected amplicon size) in qPCR with both F1/R0 and F1/R1 primers/probe pairs indicating amplification specificity (Figure S2B). Based on these results, we proceeded to use F1/R1 primers in the qPCR assay together with the TaqMan probe for detection of *S. aureus* in this paper.

**Table S4.** Survey questions for user feedback

| | |
|---|---|
| Was the CandyCollect easy to hold on to the handle? | (1 Very Bad - 5 Very Good) |
| Was the CandyCollect easy to suck on? | 1 - 5 |
| Does the CandyCollect look appealing (the color, the overall appearance)? | 1 - 5 |
| Did you like the taste of the CandyCollect? | 1 - 5 |
| Do you have a dry mouth today? | Yes No |
| Do you have a chronic condition such as Sjogren's syndrome that affects saliva production? | Yes No |
| If so, did you think your saliva production decreased during your participation in this study? | Yes No |
| Would you recommend these CandyCollect to children ages 4 and above? | Yes No |
| Please explain your choice from the previous question. | |
| How can we improve our CandyCollect for children? (Provide any suggestions) (optional) | |
| What was your preferred method of sampling? | CandyCollect<br>ESwab<br>Spitting Tube |
| What was the most suitable sampling method for children? | CandyCollect<br>ESwab<br>Spitting Tube |
| Please explain your choice from the question above | |
| What method provided the best sampling experience? | CandyCollect<br>ESwab<br>Spitting Tube |
| Rank the methods from easiest to hardest to use. | CandyCollect<br>ESwab<br>Spitting Tube |
| What method provided the best sampling experience? | CandyCollect<br>ESwab<br>Spitting Tube |
| Which method seemed the least invasive? | CandyCollect<br>ESwab<br>Spitting Tube |
| Which method was the least disgusting or uncomfortable? | Yes No |
| Which method was the most sanitary? | |
| Was it easy to follow the instructions? | 1 – 5 |
| How easy was it to complete the surveys and enroll in our study? | 1 – 5 |
| How can we improve our instructions and the survey section of this form? (Provide any suggestions) (optional) | |

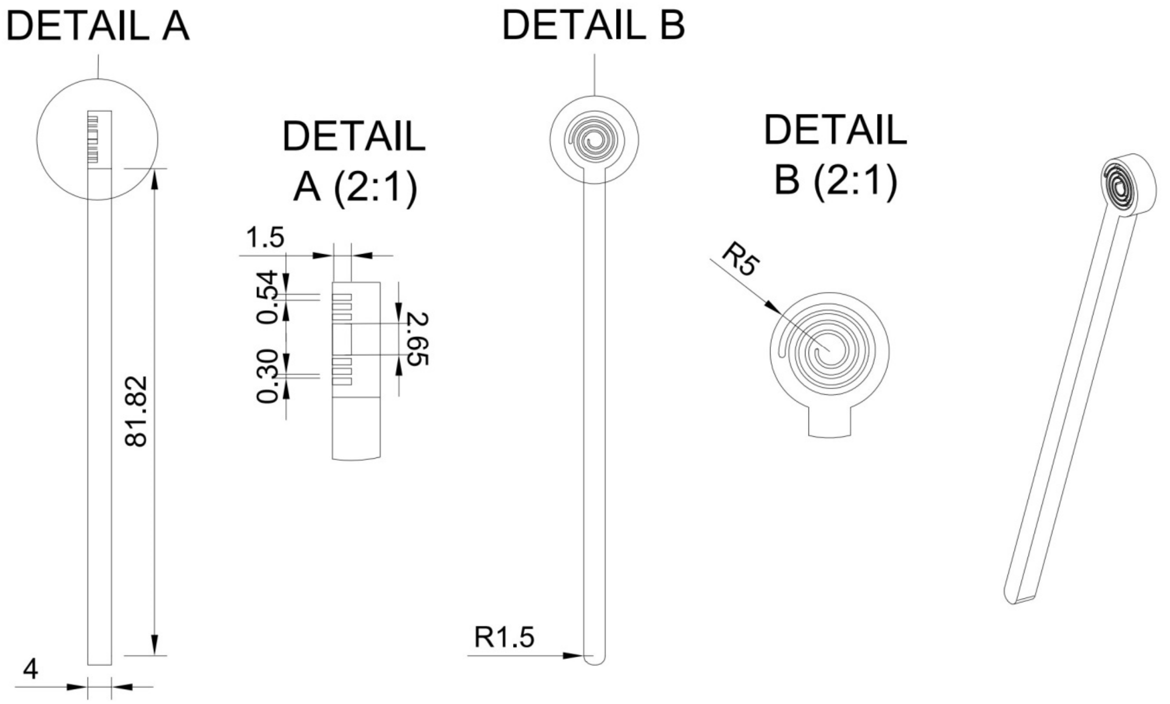


**Figure S1**. Schematic of the CandyCollect device dimensions. All dimensions are in mm.

qPCR amplification curves and standard curves with SYBR green chemistry

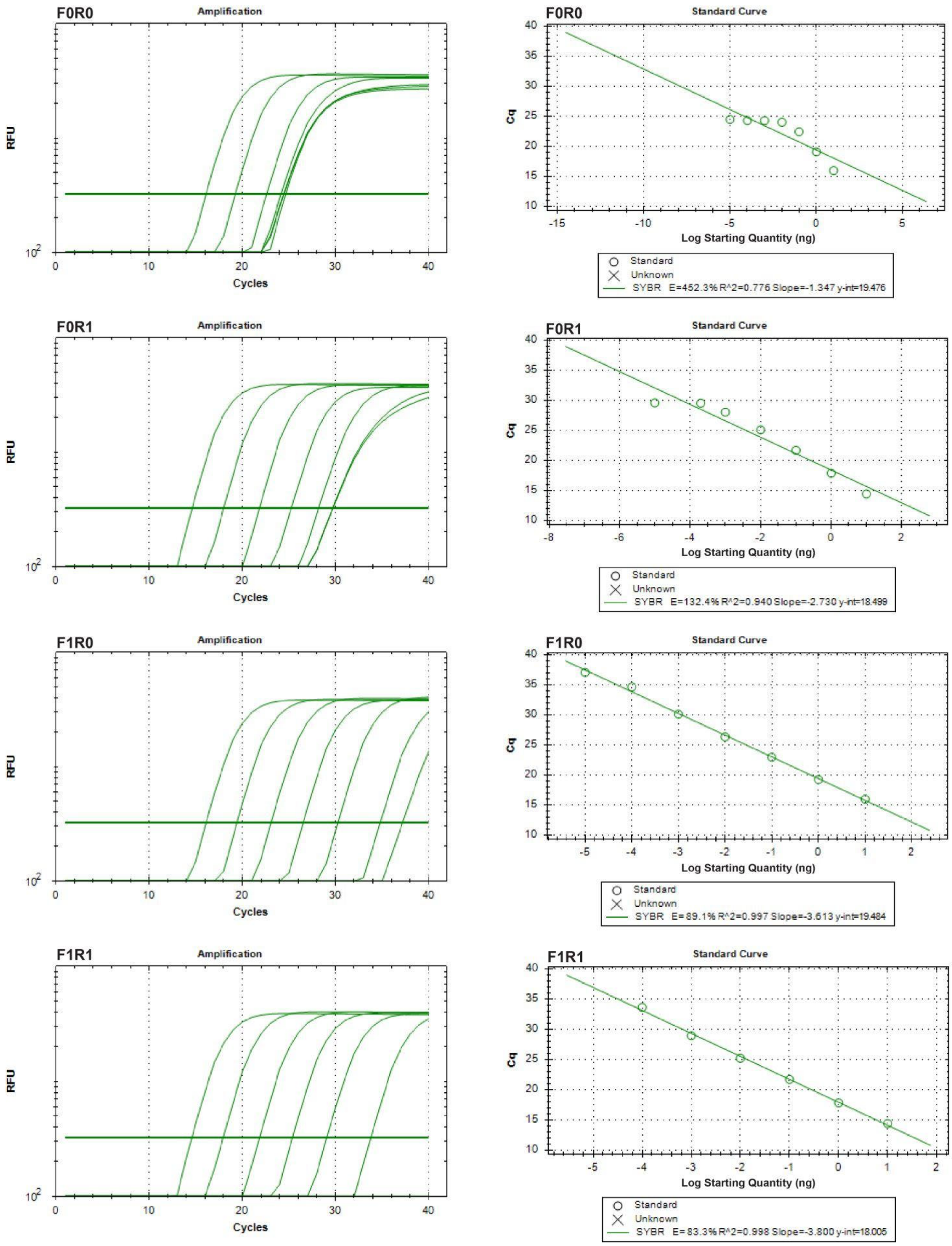


**Figure S2**. *S. aureus* primer validation by qPCR with SYBR Green chemistry. The qPCR amplification curves and standard curves were from different combinations of *S. aureus* forward and reverse primers. SYBR Green chemistry was used for detection. F0 and R0 are the original forward and reverse primers used in Galia et al. 2019[1]; F1 and R1 are modified forward and reverse primers, respectively.

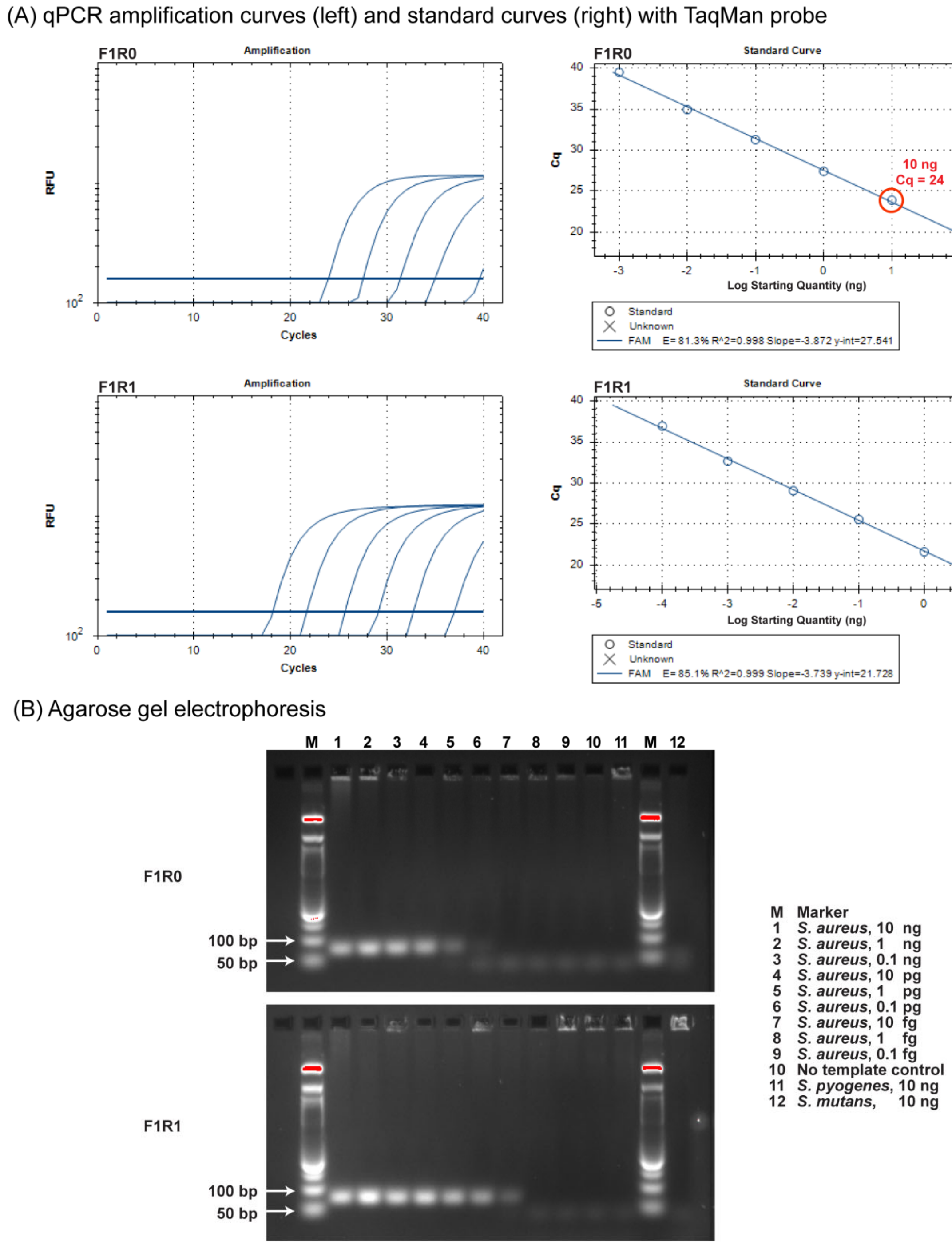


**Figure S3**. *S. aureus* primer validation by agarose gel electrophoresis and qPCR with TaqMan probe. The modified forward and reverse primers (F1/R1) with probe analysis used in this paper for *S. aureus* detection. (A) The qPCR amplification curves (left) and standard curves (right) of *S. aureus.* Probe was added in the qPCR assay. The result showed that the F1/R1 pair (bottom) had better sensitivity and wider dynamic range compared with F1/R0 pair (top). (B) Agarose gel electrophoresis demonstrated high selectivity of the qPCR assay. Agarose gel electrophoresis showed a single PCR product (~70 bp, expected amplicon size) in qPCR with both F1/R0 and F1/R1 from primers/probe pairs (top: F1R0 and bottom: F1R1) indicating amplification specificity. The templates were 1:10 serial dilution of DNA (10 ng to 0.1 fg /reaction) from *S. aureus* DNA (lane M). No PCR products were shown in DNA samples from no template control (lane 10), *S. pyogenes* (lane 11) (10 ng), and *S. mutans* (lane 12) (10 ng). 3% agarose gel was used to separate DNA products from the qPCR reactions.

(A) *S. mutans* standard curves for human participant samples

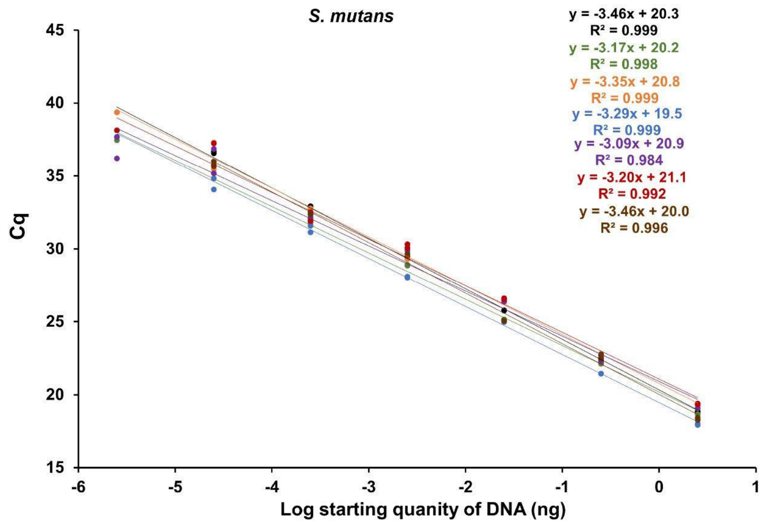


(B) *S. aureus* standard curves for human participant samples

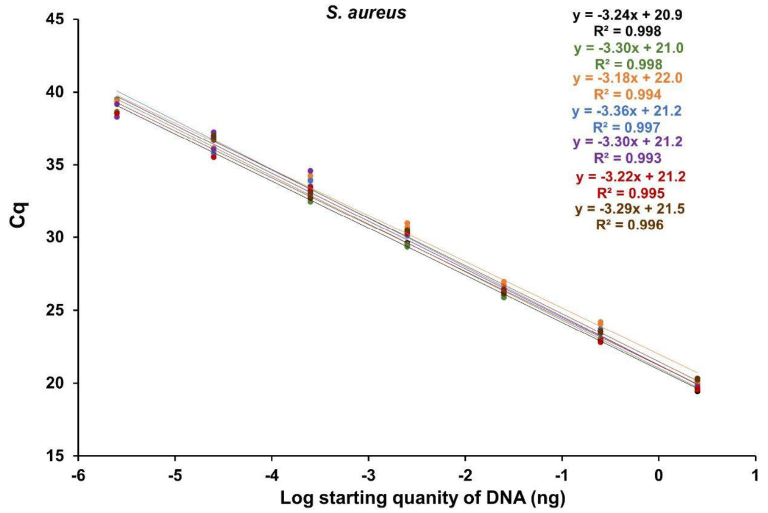


**Figure S4.** Standard curves for the (A) *S. mutans* and (B) *S. aureus* qPCR assays. 1:10 serial dilutions of genomic DNA ranging from 25 ng to 25 fg were used as template for qPCR. Each dot represents one technical duplicate (in cases where one point is visible the duplicates were identical). The standard curves in which Cq values were plotted against starting template DNA, were linear. qPCR slopes ranged from -3.09 to -3.46 for *S. mutans* and -3.18 to -3.36 for *S. aureus* across 7 independent experiments.

(A)

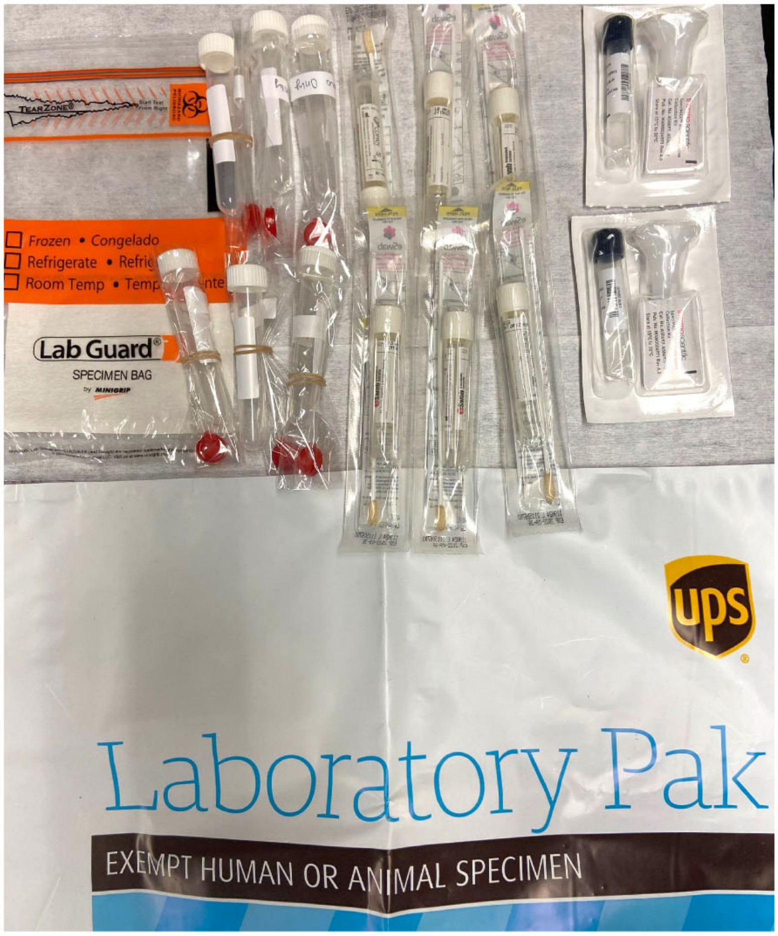


(B)

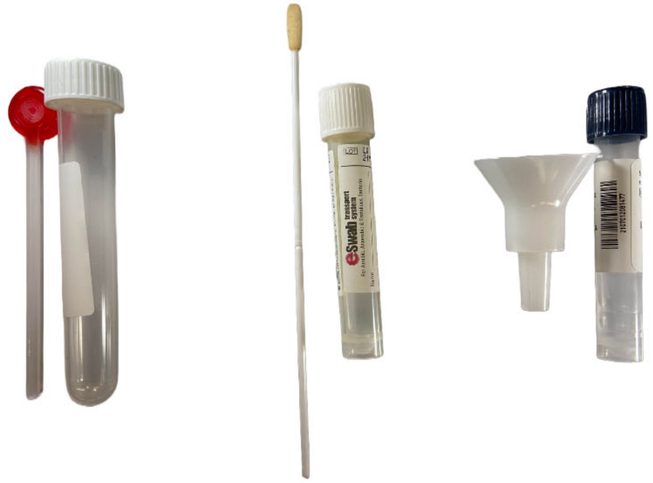

**Figure S5**. At-home saliva sampling devices for human subjects study. Self-saliva collection methods enable at-home sampling then shipping back to lab analysis. Three different collection methods included six CandyCollect devices, six ESwab™, and two SpeciMAX Stabilized Saliva Collection Kits (from left to right). (A) wrapped (B) unwrapped.

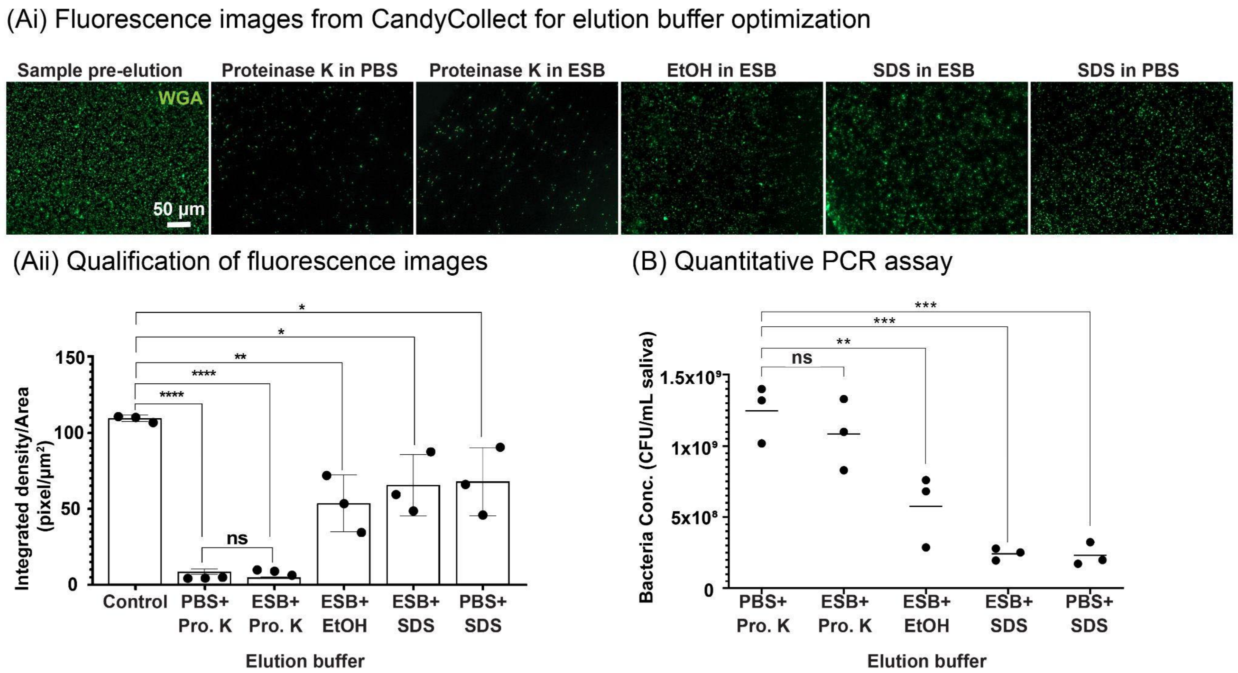


**Figure S6.** Additional elution experiments demonstrate that *S. aureus* captured by CandyCollect devices can be removed efficiently via elution buffers. (A) *S. aureus* at a concentration of $1x10^9$ CFU/mL was incubated on the CandyCollect device and eluted via five elution buffers. The image result suggests that only the Proteinase K in PBS and Proteinase K in ESwab buffer (ESB) could efficiently remove *S. aureus* on CandyCollect. *S. aureus* was green fluorescently labeled with WGA. (Aii) Quantification of the integrated density per area (pixel/$\mu m^2$). Each data point represents an individual CandyCollect; The bar graph represents the mean ± SEM of n = 3 CandyCollects. Data sets were analyzed using one-way ANOVA; p-values are indicated for pairwise comparisons between the control and different elution buffers:$*p \leq 0.1$, $**p \leq 0.01$, $****p \leq 0.0001$ (Tukey's multiple comparison tests). (B) Proteinase K in PBS and Proteinase K in ESwab buffer were the most efficient elution buffers based on the qPCR results. Quantification of *S. aureus* by qPCR. Each data point represents an individual CandyCollect. No significant differences were observed between Proteinase K in PBS and Proteinase K in ESwab buffer.

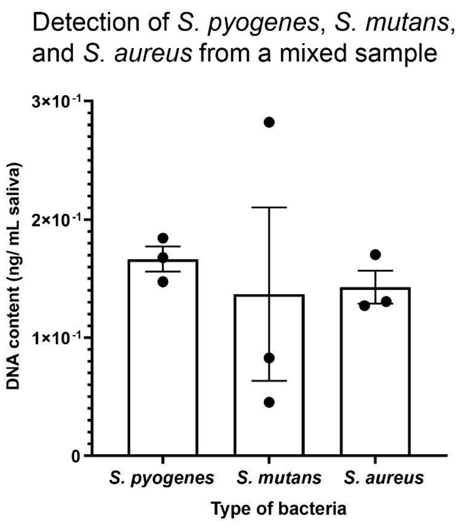


**Figure S7.** qPCR tests demonstrate that the mixture of three bacteria in saliva can be analyzed for their individual concentrations. Three bacteria, *S. pyogenes*, *S. mutans*, and *S. aureus*, were mixed at the concentration of $10^4$ CFU/mL. Quantification of the three bacteria by qPCR. DNA contents were detected in a bacterial concentration-dependent manner. Each data point represents an individual CandyCollect; the bars represent the mean ± SEM of n=3 CandyCollects.

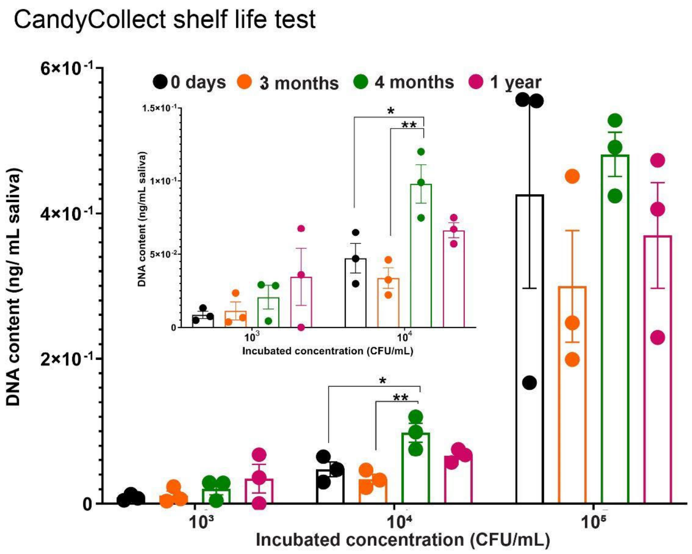


**Figure S8.** qPCR shelf life tests demonstrate that the CandyCollect device effectively captures *S. pyogenes* after 1 year of storage. CandyCollect devices were plasma treated and stored at room temperature for 0 days (control group), 3 months, 4 months, and 1 year. After the storage period, *S. pyogenes* was incubated on the CandyCollect devices, eluted, and analyzed by qPCR. Each data point represents an individual CandyCollect device; the bar graph represents the mean ± SEM of n = 3 CandyCollect devices. Data sets were analyzed using one-way ANOVA followed by Tukey's multiple comparisons test; p-values are indicated for pairwise comparisons between different storage times (*p<0.05 and **p<0.01). Note: one of the CandyCollect devices from 1 year shelf life $10^3$ CFU/mL had no qPCR signal.

**References**

(1) Galia, L.; Ligozzi, M.; Bertoncelli, A.; Mazzariol, A. Real-time PCR assay for detection of Staphylococcus aureus, Panton-Valentine Leucocidin and Methicillin Resistance directly from clinical samples. *AIMS Microbiol.* **2019**, 5, 138-146.
